# A periodic table of bacteria?: Mapping bacterial diversity in trait space

**DOI:** 10.1101/2025.07.11.664459

**Authors:** Michael Hoffert, Evan Gorman, Manuel E. Lladser, Noah Fierer

**Affiliations:** Department of Ecology and Evolutionary Biology, University of Colorado Boulder, Boulder, CO USA; Cooperative Institute for Research in Environmental Sciences, University of Colorado Boulder, Boulder, CO USA; Department of Applied Mathematics, University of Colorado Boulder, Boulder, CO USA

## Abstract

Bacterial diversity can be overwhelming. There is an ever-expanding number of bacterial taxa being discovered, but many of these taxa remain uncharacterized with unknown traits and environmental preferences. This diversity makes it challenging to interpret ecological patterns in microbiomes and understand why individual taxa, or assemblages, may vary across space and time. While we can use information from the rapidly growing databases of bacterial genomes to infer traits, we still need an approach to organize what we know, or think we know, about bacterial taxa to match taxonomic and phylogenetic information to trait inferences. Inspired by the periodic table of the elements, we have constructed a ‘periodic table’ of bacterial taxa to organize and visualize monophyletic groups of bacteria based on the distributions of key traits predicted from genomic data. By analyzing 50,745 genomes across 31 bacterial phyla, we used the Haar-like wavelet transformation, a model-free transformation of trait data, to identify clades of bacteria which are nearly uniform with respect to six selected traits - oxygen tolerance, autotrophy, chlorophototrophy, maximum potential growth rate, GC content and genome size. The identified functionally uniform clades of bacteria are presented in a concise ‘periodic table’-like format to facilitate identification and exploration of bacterial lineages in trait space. While our approach could be improved and expanded in the future, we demonstrate its utility for integrating phylogenetic information with genome-derived trait values to improve our understanding of the bacterial diversity found in environmental and host-associated microbiomes.

## Introduction

The periodic table of elements offers an interpretable framework for organizing atomic elements based on their properties [1]. This framework has been adapted across various fields to categorize the equivalent of elements, from food types to cell types, according to shared properties [2–5]. In fields like microbial ecology where organisms are the subjects of study, microbial taxa are documented in databases like the Genome Taxonomy Database [6] and BacDive [7] which serve to organize reference genomes, phylogenetic trees, and trait information. However, these databases omit a key feature of a periodic table: organizing units according to functional information. Microbial traits are as much the central focus of microbial ecology as chemical properties are the focus of physical chemistry: traits of microbes are analogous to chemical properties of atoms, where each determine the outcomes of small-scale interactions in their respective systems. Therefore, building a periodic table of microbial diversity that clearly organizes taxa and lineages based on shared traits would highlight key differences and illustrate patterns or periodicity in what we know, or think we know, about the diversity of bacterial life on Earth. Although applying this idea has been considered in some biological contexts (e.g. ecological niches in Winemiller et al., 2015), there is no consensus about how to interpretably measure, organize, or visualize the distributions of microbial traits across large swaths of taxonomic diversity. Establishing a framework for associating bacterial taxonomic identify with trait distributions would not only enhance our understanding of microbial ecology but also facilitate meaningful comparisons and insights across the rapidly expanding landscape of microbial research.

Despite its promise, constructing a “periodic table of bacterial taxa” (PTBT) which succinctly visualizes bacterial trait-taxonomy relationships is challenging. Finding and assigning trait measurements to taxa in the PTBT is hampered by the intrinsic difficulty of measuring most bacterial traits, nebulous or dated taxonomic assignments [9, 10], or complex trait distributions arising from horizontal gene transfer [11] and variable rates of evolution [12]. Although commonly-cited examples of broad taxonomic groups with conserved metabolisms exist, such as photoautotrophic *Cyanobacteria* [13], most bacterial traits are “distributed in phylogenetic clusters with a continuum of depths” [14] and cannot be uniformly attributed to a taxonomic group without empirical evidence. Establishing such empirical estimates of trait values for taxonomic groups is an evolving challenge. The traits of many bacterial taxa remain unknown because they are resistant to cultivation and not readily amenable to experimentation. However, cultivation-independent methods which infer trait values from genomic data [15] promise to circumvent these challenges and make it feasible to infer some traits from genomic data alone. As ongoing sequencing efforts yield thousands of new bacterial genomes per year, a combination of cultivation-independent and dependent methods have yielded databases of bacterial traits for an increasing diversity of bacteria (see Barberán et al., 2017; Madin et al., 2020; Reimer et al., 2019). A wide range of bacterial traits can now be inferred from genomic information, including autotrophy [18–20], nitrogen fixation [21], antibiotic resistance [22], maximum potential growth rates [23], temperature tolerances [24–26], pH preferences [27], and oxygen tolerances [28]. These advances have provided new means by which traits can be inferred for uncultivated and cultivated taxa alike, providing the empirical estimates of traits needed to build a PTBT.

Here we show how a PTBT can be constructed using recently developed approaches to identify and visualize phylogenetic groups with particular trait distributions. Our approach addresses the challenges of finding trait information for most bacteria and interpreting the complex distribution of those traits in large phylogenies in two parts; first using genome-derived trait estimates to characterize a large swath of bacterial phylogenetic diversity and then applying the recently developed Haar-like wavelet projection (HWP) [29] to identify phylogenetic groups with conserved traits (Figure 1). Here we focus on six key traits (genome size, GC content, oxygen tolerance, autotrophy, chlorophototrophy, and maximum potential growth rate), estimating each trait for over 50,000 representative genomes in the Genome Taxonomy Database (GTDB, Parks et al., 2018). We use the HWP to compute the trait variance associated with each phylogenetic node and identify clades with minimal trait variation - clades which can be accurately represented in single cells in a prototype periodic table (Figure 1A,B). Because many traits exhibit some degree of phylogenetic conservation [14, 30], we expect that a small number of phylogenetic splits can capture large amounts of trait variance, and identification of these splits via HWP will make it feasible to design a PTBT simple enough to link taxa to traits while also accurately describing most of the variance in these traits (Figure 1C-D). Our approach represents an interpretable and quantifiable method to systematically characterize the largest sources of trait variance across the bacterial tree of life. Our use of quantitative means to discover and arrange the cells of the PTBT is deliberate, as we expect that this method can ultimately be expanded to add additional traits, improve inferences of individual trait values, and include additional taxa as genome-based models, experimental training data, and genomic datasets continue to improve.

**Figure 1.**
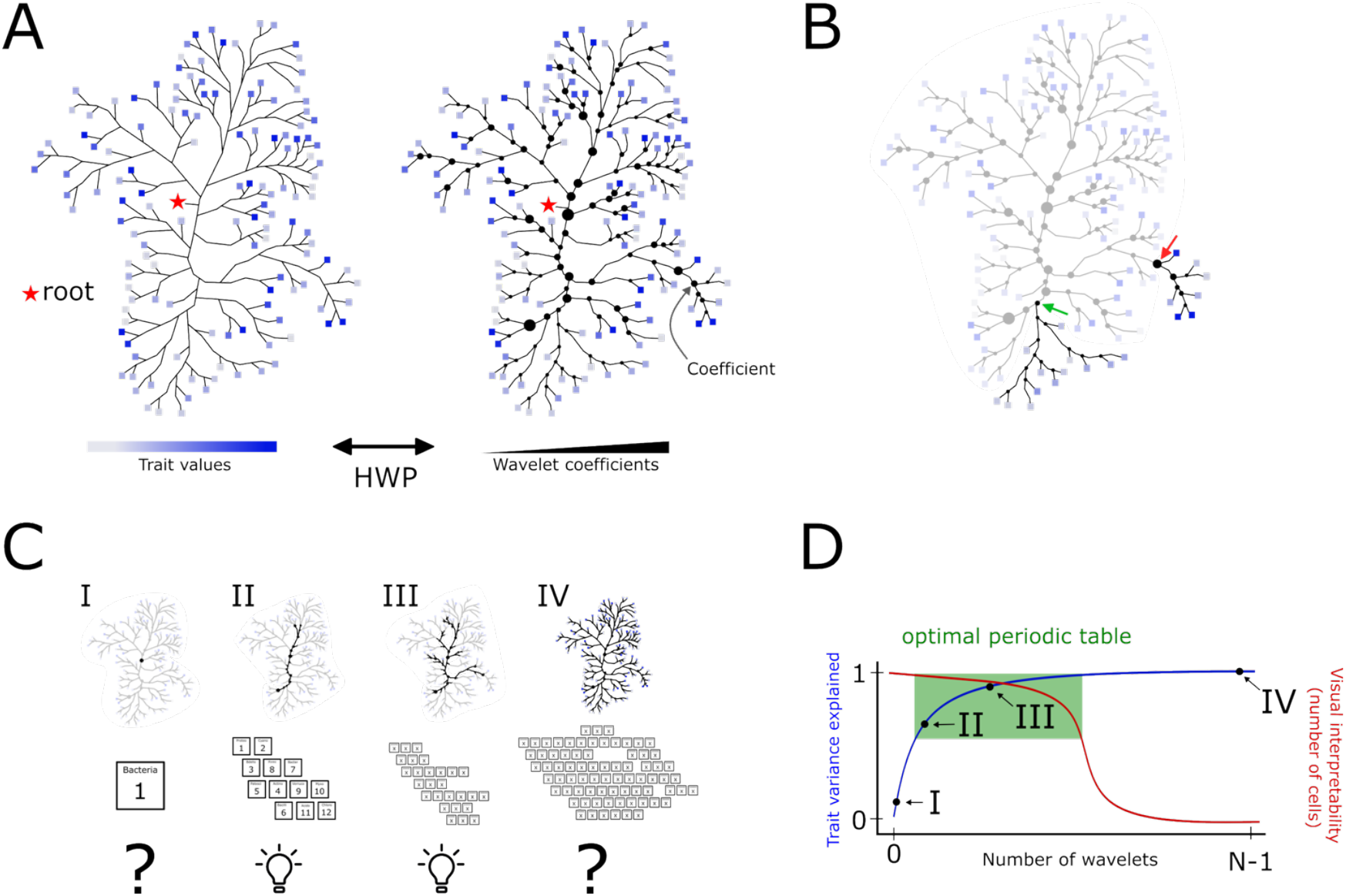
Conceptual outline of our method for constructing a periodic table of bacterial taxa (PTBT). The goal of a PTBT is to describe the distribution of traits across bacterial taxa for phylogenies with hundreds of thousands of leaves using quantitative methods. (A) We first apply the Haar-like wavelet projection (HWP) to a large, complex phylogeny with a trait measured for each leaf, e.g. a trait value predicted from a bacterial genome. The resulting Haar-like wavelet coefficients are associated with internal nodes of the phylogeny and measure the amount of variance uniquely associated with each node. (B) Our method of constructing a PTBT identifies the ancestral nodes of clades with small wavelet coefficients (green arrow), which are monophyletic groups with relatively uniform trait values. In contrast, clades with large wavelets (red arrow) cannot be accurately summarized by representing the collection of leaves with their average trait values. (C) The PTBT is constructed by collapsing clades with minimal variance in the trait to their ancestral node and using the tips of the collapsed phylogeny as “cells” in the table. The degree of collapsing is continuous: complete collapsing of the phylogeny (resulting in a single cell, tree I) is undesirable, as is the original, complex phylogeny (tree IV). An intermediate representation (trees II and III) illustrates which clades are functionally variable or uniform in a visually interpretable number of cells. (D) Therefore, the optimal PTBT selects the X largest wavelets to summarize trait-taxon relationships while preserving visual interpretability.

## Results

### Compilation of trait data for 50,745 bacterial genomes

To accomplish the central task of arranging bacterial taxonomic in trait space, we first downloaded 62,291 species-level representative genomes from the Genome Taxonomy Database (GTDB) version 207 [6] for trait estimation. We used representative genomes only to ensure most genomes were relatively complete, uncontaminated, and had a standardized taxonomic assignment aligned with the GTDB phylogenetic tree. Our analyses were restricted to the 31 bacterial phyla which included at least 100 species-level representative genomes per phylum and only genomes with at least one ribosomal protein, a requirement for growth rate predictions [23], yielding 50,745 of the original 62,291 bacterial genomes. For each of these 50,745 genomes, we inferred values for six ecologically relevant traits: oxygen tolerance, autotrophy, chlorophototrophy, maximum potential growth rate, GC content, and genome size as described below and in the Methods section which includes full details on how these traits were determined from the genomic data. Supplementary Table 1 includes the inferred trait values for all 50,745 genomes. We emphasize that our trait inferences, particularly the inferences of autotrophy, phototrophy, and O_2_ tolerance, are approximations which do not include all possible pathways for these functions and could ultimately be improved. However, as detailed below and in Figure 2, our inferences broadly align with published literature, despite the literature’s bias toward taxa that are readily cultured and studied *in vitro*. Likewise, we describe the general patterns at broad taxonomic levels, acknowledging that the phylogenetic depth at which taxa exhibit consistency in traits varies depending on the trait and lineage in question (as discussed in more detail in the following sections).

**Figure 2.**
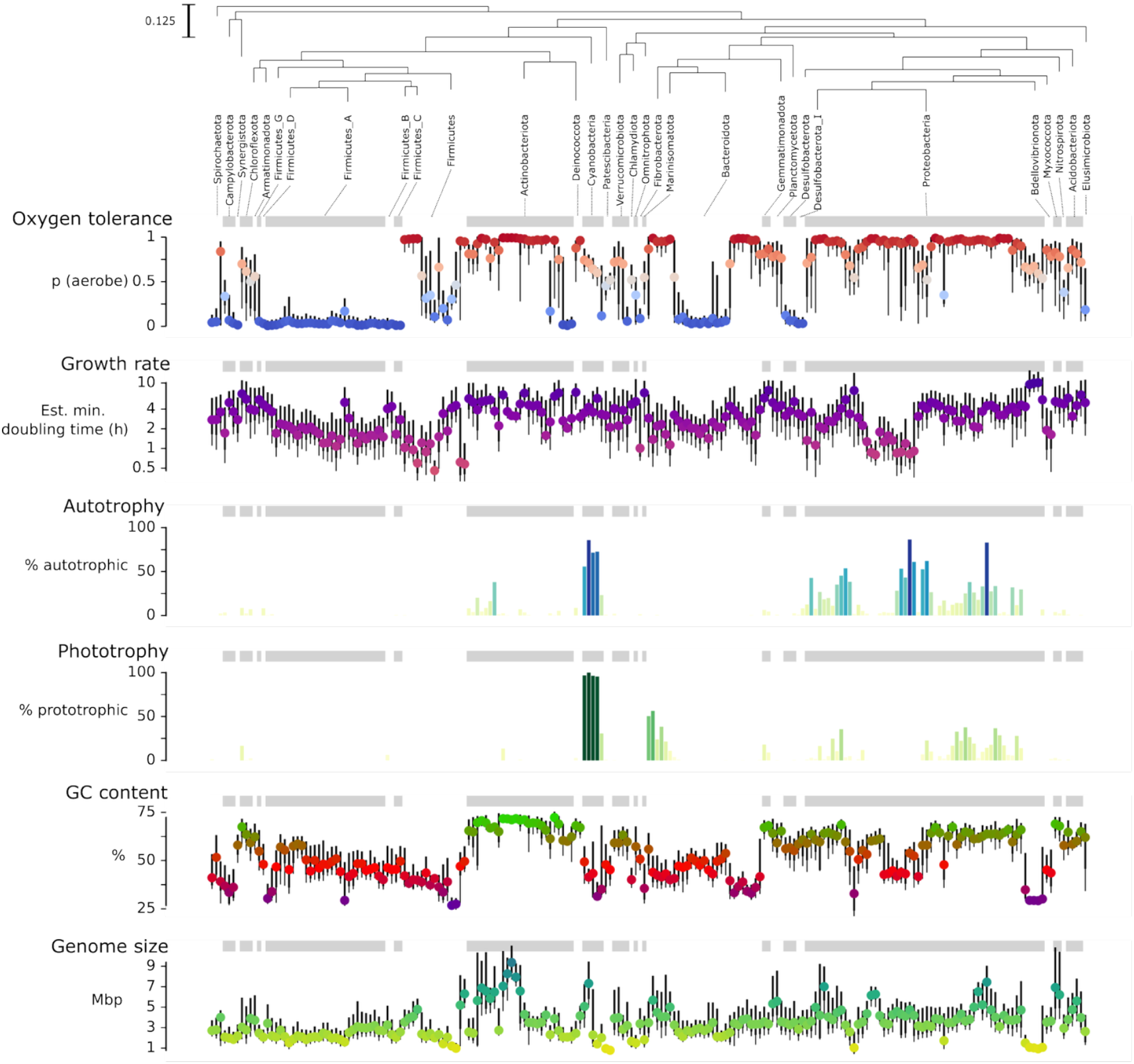
Distributions of estimated trait values for 50,745 GTDB representative genomes for six key traits. To ensure the data from all of these phyla are visible on this plot, the data are collapsed into bins represented by boxplots: each phylum is represented by at least one boxplot, where each boxplot shows a color-coded median and the 10^th^,25^th^, 75^th^, and 90^th^ quantile values for each trait across up to 500 genomes. Gray bands at the top of each plot act as a visual aid to delimit the boxes corresponding to each phylum using the phylogenetic placement of 31 bacterial phyla from GTDB v207 (Parks et al. 2022). Because autotrophy and phototrophy are binary variables, they are shown as bars representing the fraction of genomes in each bin that are positive for that trait. The six measured traits are shown in the following order: (1) oxygen tolerance, the probably of being aerobic, predicted using a random forest trained on BacDive genomes (2) maximum potential growth rate in estimated minimal doubling time, estimated with gRodon, (3) phototrophy, predicted using the presence of bacteriochlorophyll synthesis genes, (4) carbon fixation, predicted using the presence of key genes, (5) genome GC content and (6) genome size computed directly from genomic data. See Methods for details on how these traits were inferred.

GC content and genome size were calculated directly from genomic data and included because these properties are associated with ecological attributes, from environmental tolerances to oxygen, light and temperature [31–34] to specific metabolisms [35] and ecological strategies [36, 37]. Both genome size and GC content were considered as continuous trait values ranging from 0.22 – 25 Mbp and 15 -77%, respectively, across the 50,745 bacterial genomes. The taxa with very large genomes included many Actinobacteriota, Myxococcota, and specific clades of Proteobacteria and Cyanobacteria, while those with smaller genomes were found across many phyla - most Firmicutes, some Proteobacteria, and phyla like Omnitrophota and Patescibacteria where small genome sizes are a distinctive trait [37, 38]. The genomes with the highest GC content were common in Actinobacteriota, Proteobacteria, and sister clades, while fewer and more specific subgroups of Firmicutes, Proteobacteria, and Bacteroidota contained low-GC genomes (Figure 2 and Supplementary Table 1).

Maximum potential growth rate was included because it is a key feature distinguishing variation in general life history strategies across bacteria [39, 40] with maximum potential growth rate approximated using minimum potential doubling time predictions from gRodon v2 [41]. Values for maximum potential growth rate ranged from 0.01 to 25 hours, with lower values indicating faster potential growth and shorter generation times. Taxa with low estimated generation times (<1 hour) were relatively uncommon but phylogenetically widespread across anaerobic Firmicutes and aerobic Proteobacteria. Long estimated generation times (slowest maximum potential growth) were prevalent in many groups, including specific Proteobacteria, many Actinobacteriota, and Gemmatimonadota (Figure 2). Taxa that have been isolated *in vitro* almost always have shorter predicted generation times (faster maximum potential growth rates) than corresponding, uncultivated taxa within the same phylum from which genomes were obtained via assembly from metagenomes (Supplementary Figure 8), highlighting a bias for faster growth among taxa that are readily cultivable [23].

The potential for autotrophic metabolism, or the ability to fix CO_2_, was estimated using key enzymes and pathway completeness [18, 42] from two of the four known bacterial autotrophy pathways (Calvin Cycle and hydroxypropionate bi-cycle) which are reasonably well-characterized. This trait is encoded as a binary variable (either inferred to be capable or incapable of autotrophic CO_2_ fixation) when a genome contained sufficient completeness of either pathway. Using this approach, 9.3% (4,627) of the bacterial genomes were inferred to be capable of autotrophic metabolism via either of the two measured pathways. As expected, the inferred capacity for autotrophic CO_2_ fixation was most prevalent among Cyanobacteria and Proteobacteria, and otherwise found in small numbers of taxa in Actinobacteriota [43] sister phyla of the Firmicutes supergroup [44–47] (Figure 2).

Similarly, we screened all genomes for key genes and pathways to infer chlorophototrophy, the capacity to capture light energy using chlorophyll. Approximately 6% (3,223) of bacterial genomes contained chlorophyll synthesis genes ubiquitous in a genomic dataset of chlorophototrophs [20]. Consistent with the literature [13, 20, 48–50], the inferred capacity for chlorophototrophy was largely (but not always) restricted to taxa within the phyla Proteobacteria, Cyanobacteria, Chloroflexota, and other groups which were also inferred to be autotrophic (Figure 2)

Finally, oxygen tolerance was predicted from gene presence using a random forest model trained on characterized taxa from BacDive [7]. Oxygen tolerance was encoded as the probability each genome is aerobic (zero to one) and was included because tolerance of oxygen is a key determinant of metabolic strategies [51] and the potential environments in which taxa can thrive. As expected, oxygen tolerance is not strongly conserved at the phylum level (Figure 2). Aerobes are particularly prevalent in Proteobacteria, Actinobacteriota, and at least some clades of most other phyla, while taxa within Firmicutes A, Desulfobacterota, and subgroups of some Bacteroidota were almost exclusively inferred to be anaerobes.

### Identification of taxa with shared traits using Haar-like wavelets

To identify lineages with conserved traits across all 50,745 bacterial genomes (from the genome-derived inferences described above), we summarized trait-phylogeny relationships using the Haar-like wavelet projection (HWP). The HWP performs phylogenetically-informed analyses of trait values associated with species, or leaves, into measurements of the trait variation uniquely attributed to each interior node of the phylogeny (see Gorman & Lladser, 2024 for specific methods). The magnitude of a “wavelet coefficient” associated with each node quantifies the degree of change observed between the leaves descended from any given node in the phylogenetic tree. The wavelet coefficients normalized to fraction of total variance are shown in Supplementary Figure 2 and reveal that most trait variance is explained by only a few internal nodes. For example, at least 25% of the variation in oxygen tolerance, growth rate, GC content, and genome size are attributed to one split (clade c000130) at the common ancestor of slow-growing high GC aerobes with large genomes (Actinobacteriota) and fast-growing low-GC anaerobes with small genomes (most Firmicutes). Fewer than 100 wavelets are required to explain 60% or more of total trait variance (Figure 3); 26 of these wavelets are shared across two or more traits, implying that trait changes frequently occur at the same phylogenetic nodes, perhaps due to coordinated evolution of traits [14]. Additionally, many of the phylogenetic splits with large wavelet coefficients reside at deeper nodes, often at approximately the phylum level of resolution (Supplementary Figure 3). Collectively the HWP reveals that these six traits are correlated, significant sources of variation are found deep in the phylogeny, and small numbers of splits can describe trait distributions accurately, all observations which indicate that a PTBT can describe a majority of the trait variance across a broad diversity of bacteria, even when considering multiple traits simultaneously.

**Figure 3.**
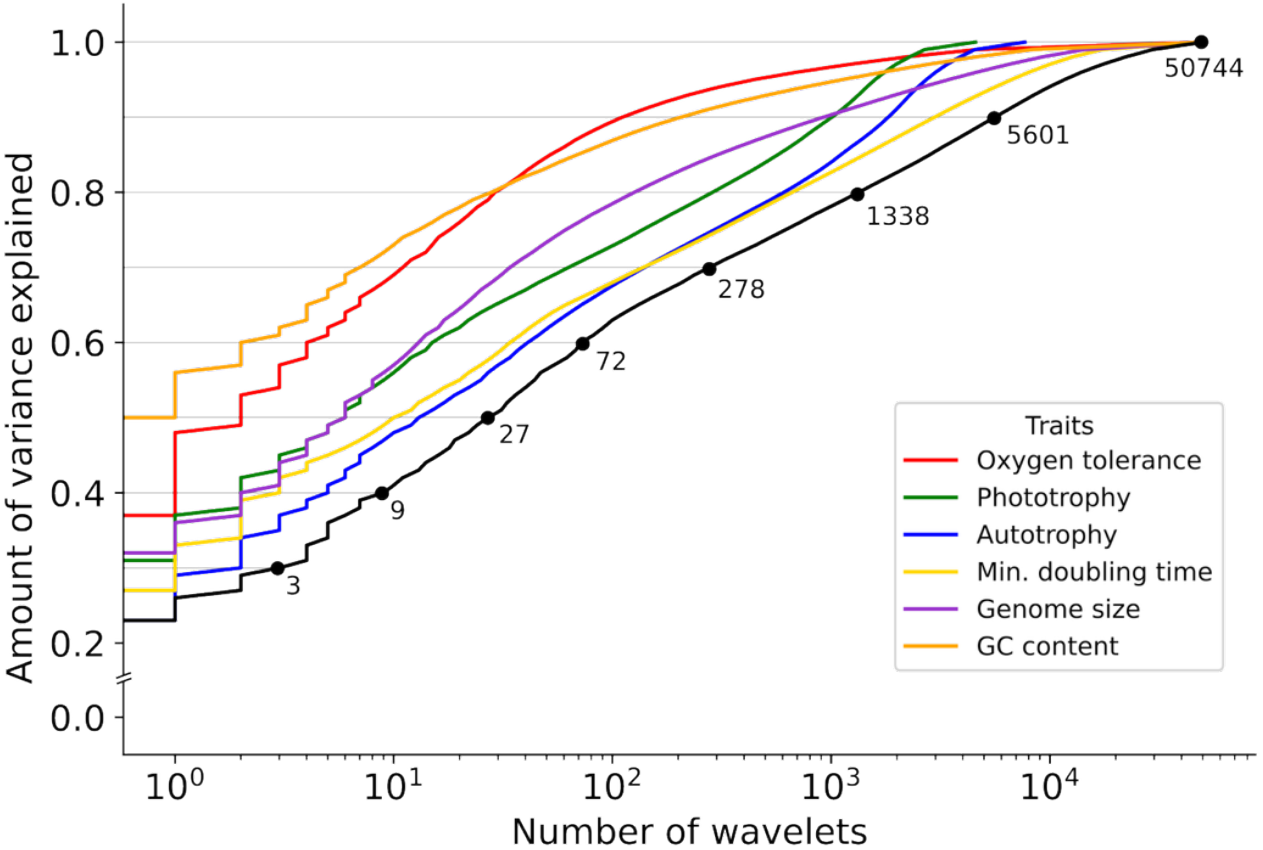
Per-coefficient variance explained for each trait versus the number of wavelets used to describe that variance. Each curve is constructed by sorting the wavelet coefficients for each trait by magnitude and summing the magnitudes. The intersection of horizontal bars at 10% variance intervals indicate the total number of unique wavelet coefficients (or phylogenetic splits) which explain that degree of total variance for the trait. For the periodic table, we selected a threshold of 60% variance explained, which yielded a total of 1307 ‘cells’, or groupings of bacteria, with the 256 cells which contained the largest number of unique genomes (∼80% out of the 50,745 genomes) shown in Figure 5, our ‘periodic table’.

Association of traits with one another and phylogenetic structure is a desirable condition given that the utility of the PTBT is tied to its visual complexity and information fidelity. The deconvolution of phylogeny-trait relationships provided by the HWP illustrate that trait distributions and co-occurrence are not combinatorially complex, and that trait-taxon relationships can be simplified while preserving variation in a PTBT. Other studies have also determined that vertical transmission and deep conservation are properties of complex bacterial traits [14, 30], but our data reveal that these trends are not expressed equally among the traits examined here. The small number of highly explanatory wavelets (Figure 3 and SF2) in the autotrophy and growth rate traits, and to lesser extents in phototrophy and genome size, indicate these traits have relatively lower degrees of phylogenetic conservation (and may be more difficult to summarize). The different degrees of conservation in genome size and GC content, measured directly from genomes, indicate that variation in phylogenetic conservation is biological and not simply due to inference errors. Among our traits, various explanations may explain varying degrees of phylogenetic association, from a history of horizontal transmission in the Calvin Cycle [18], to selection or drift altering genome size based on the specific biotic and abiotic factors present [36, 52–54]. Such variations make visual properties which depict trait co-occurrence and depth of phylogenetic conservation important components of the PTBT’s visual design.

### Drafting a ‘periodic table’ to visualize bacterial diversity in trait space

The procedure for constructing the PTBT involves analyzing the wavelet coefficients to find parents of sister clades whose leaves *are not similar* with respect to each trait and therefore do not represent the functionally uniform (or nearly uniform) clades that we want the cells of the PTBT represent. At least 60% of trait variance can be explained by only 72 unique wavelet coefficients and corresponding nodes (Figure 3). The PTBT is constructed to explain a particular amount of variance in the data: the amount of trait variance explained (TVE) is arbitrary but determines the visual complexity of the PTBT, so we picked a value near the inflection point of the wavelet magnitude versus variance explained distribution, maximizing TVE while minimizing number of cells in the PTBT. The 72 unique nodes required to explain 60% of the variance were then used to prune branches and create a simplified phylogeny containing junctions which contribute to TVE. This pruned tree’s new leaves are clades which contain no large wavelets and therefore capture minimal descendent trait variation to the extent guaranteed by the 60% TVE threshold and are drawn in the PTBT (Figure 5). A layout of each clade in two dimensions to emulate the draft periodic table was created using each clade’s pairwise phylogenetic distance and median trait values, and computing a t-SNE-based embedding of these relationships into a common three-dimensional space with ENS-t-SNE [55]. The resulting layout combines phylogenetic information with trait information to gather cells which are similar based on phylogenetic and/or trait-based distance (Figure 4). The median trait values for each clade are drawn with different graphical elements of the cells in Figure 5A, our initial design of an empirically derived PTBT, including a diagram of the phylogeny which resulted from HWP-based collapsing to illustrate the taxonomic groups which constitute the cells in the table (Figure 5B).

**Figure 4.**
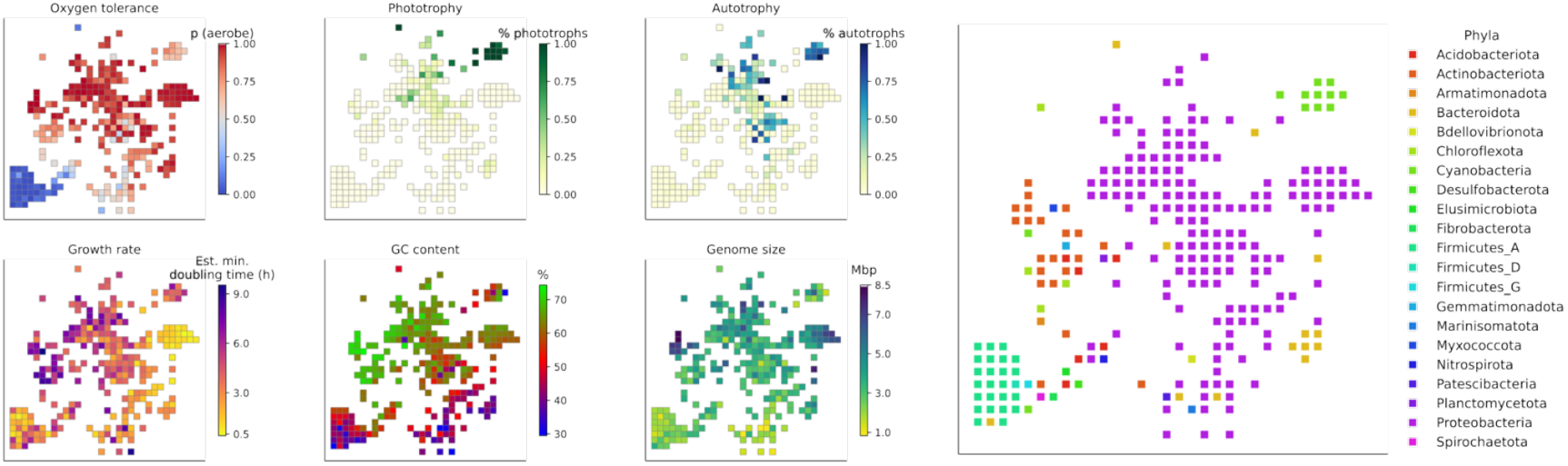
Data used to generate a layout of the periodic table diagram which combines trait and phylogenetic information using ENS-t-SNE. Each plot shows the distribution of six traits among the cells of the periodic table, with an additional plot to illustrate the phylum-level identities of the taxa within each of the cells. Each cell corresponds to a clade in the bacterial trait dataset (50,745 leaves total, 272 in this diagram) discovered by traversing the phylogeny until no sufficiently large Haar-like wavelets existed in any subtree. The median trait values from the resulting tips and a patristic distance matrix were provided to the ENS-t-SNE algorithm to generate a layout that included phylogenetic and trait information. We note that the cells and their arrangement in these panels are identical to those shown in the final periodic table shown in Figure 5, with this figure serving as a complement to Figure 5 to help visualize how traits and taxonomic identities vary across the cells in the periodic table.

**Figure 5.**
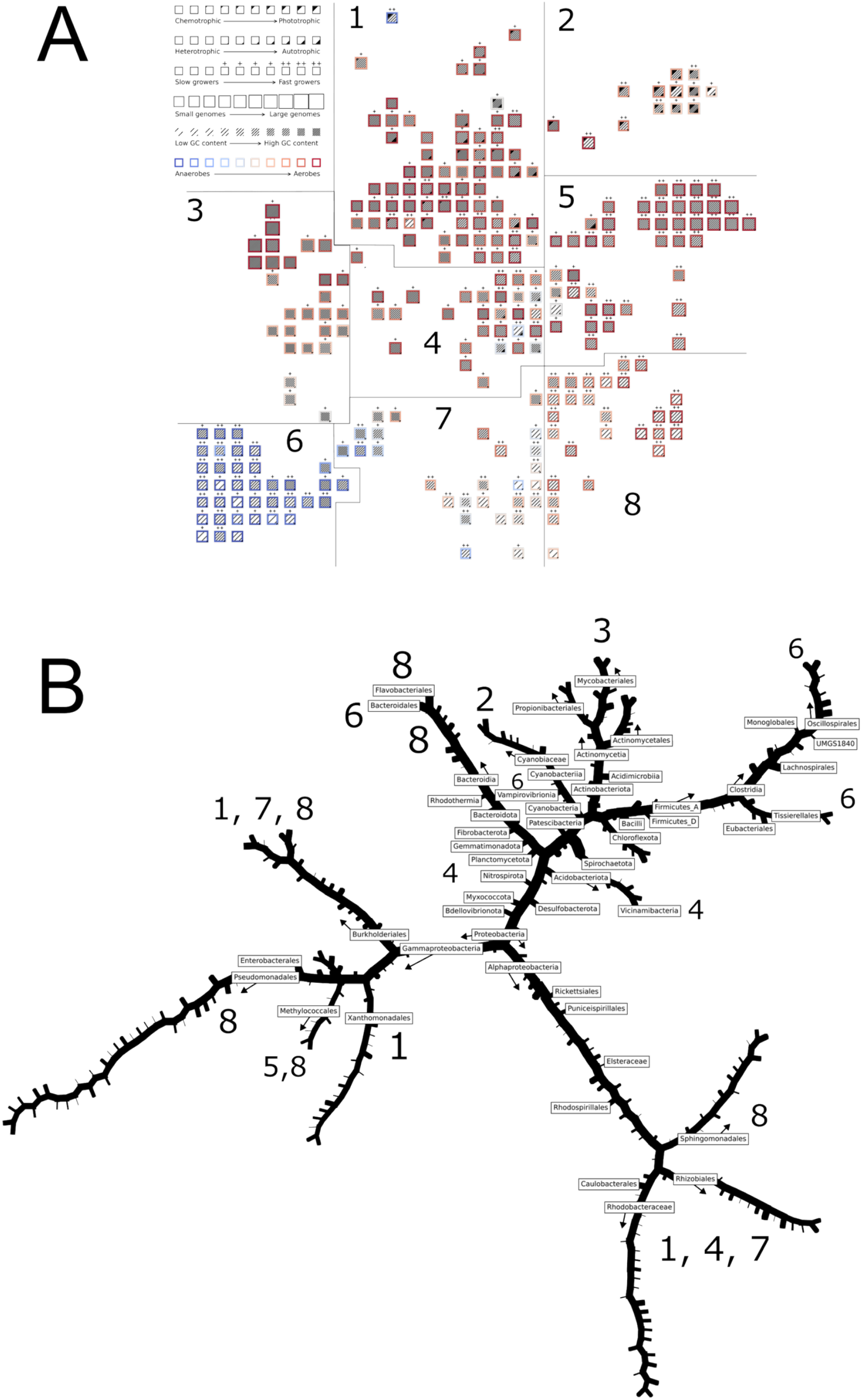
Periodic table of bacterial diversity. Functionally conserved taxonomic groups were identified by pruning the branches of a bacterial phylogeny with 50,745 leaves (species-level representative genomes) until an internal node associated with significant internal variance was discovered, essentially identifying the clades to put in a “periodic table-like” diagram. The layout of the table was generated by projecting a combination of trait and phylogenetic information into 2D space using ENS-t-SNE, then snapping points to a grid which preserved the nearby placement of clades with similar trait values and phylogenetic identity (see Figure 4). (A) A multi-trait illustration of the trait values for 272 groups of functionally homogeneous bacterial taxa discovered by the pruning algorithm, each illustrated as a single “cell.” The periodic table is subdivided into 8 parts for the purposes of description and the higher level categorization of bacterial diversity in trait space. (B) The underlying pruned phylogeny of bacteria labelling the clades which appear in the diagram from panel A (numbered, black edges), as well as some of the larger phylogenetic groups which divide the clades of bacterial taxa with homogeneous, or nearly homogeneous, trait values. The numbers indicate the regions of the periodic table (from panel A) in which some of the clades appear.

The PTBT illustrates the distributions of traits across bacterial phylogenetic diversity and facilitates the interpretation of trait co-occurrence, conservation, and divergence patterns in specific taxa. First, the PTBT illustrates the variation in the phylogenetic depths at which trait combinations are conserved. The PTBT assigns Proteobacteria, Firmicutes A, and Actinobacteriota more cells (172, 24, and 21 respectively) because traits vary more in these phyla versus Myxococcota, Cyanobacteria, Spirochaetota, Patescibacteria, and other phyla with less variation in the traits assessed here (Figure 4). Second, the layout identifies phylogenetically distinct taxa that share similar traits, separating primarily oxygen-tolerant Actinobacteriota with high GC contents and large-genomes in regions 3-4 of Figure 5A from phylogenetically distinct but otherwise similar pseudomonads in region 5 of Figure 5A. The PTBT also illustrates where related taxa have distinct traits, as in the placement of oxygen-tolerating photoautotrophic Cyanobacteriota (Figure 5A region 2) and chemoheterotrophic Bacteroidota (Figure 5A-8) that are far from each group’s single clade of chemoheterotrophic low-GC anaerobes (Bacteroidales in Bacteroidota; Vampirovibrionia in Cyanobacteria, both in region 6 of Figure 5A). Finally, the PTBT makes it possible to identify the taxonomic underpinnings of high-level associations between traits, for example in the association between GC content, oxygen tolerance, and growth rates appearing to be driven here by the distinction between Firmicutes and Actinobacteria/Proteobacteria, a result consistent with previous work [52]. Essentially, the PTBT maps trait distributions into visual channels like distance, clustering, size, and color which enables users to rapidly explore bacteria with particular groups of traits, identify the set of traits any bacterial group is likely to have, and assess the relative diversity of particular groups in trait space by examining both the number of cells in a group and their placement relative to one another.

## Discussion

This work explores how quantitative methods can help describe, explore, and visualize complex bacterial taxon-trait relationships. Inspired by the long-standing desire among microbial ecologists to find methods which identify and organize taxonomic groups with shared traits [56– 59] and recognizing that traits fundamentally determine organismal fitness and ecologies [60], we believe traits represent the most appropriate foundation on which to build a “periodic table” of bacterial diversity. Our methods of constructing a periodic table includes novel applications of genome-based trait inference methods, the HWP, and ENS-t-SNE to address persistent challenges in estimating traits across a broad diversity of bacteria and then identifying and organizing groups of bacteria into a visual framework that captures trait co-occurrence, phylogenetic conservation, and phylogenetic similarity. We can effectively organize large swaths of bacterial diversity in trait space with the PTBT because there are biological and evolutionary constraints that limit the possible combinations of trait states and allow an informative picture of bacterial taxon-trait relationships to be summarized and visualized. We expect that our approach could generate meaningful simplifications for other trait-taxon associations, with particular methods like the HWP providing interpretable information to inform the design of new representations of trait-phylogeny datasets. With adaptation, these methods could also help integrate trait data with genome databases like GTDB [6] to unify microbial functional, phylogenetic, and genomic information in a single interface.

Although efforts to synthesize traits, phylogenetic information, and visualization techniques have many realized and unrealized benefits, we acknowledge that our workflow is inherently reductive and could be improved. We have only visualized six traits with a specified degree of variance, largely ignored potential errors in trait estimation methods, and made other simplifications. In addition to improvements to trait estimations and phylogenetic inferences, whether the PTBT provides verifiable explanations of the realized patterns in ecosystems should be critically assessed in further work. In the meantime, we hope the PTBT will inspire development of new tools to summarize, assess, and validate our current knowledge - and assumptions about - the taxon-trait associations upon which many analyses in microbial ecology explicitly or implicitly rely. For example, the PTBT illustrates where assumptions can and cannot be made about the depth and variability of trait conservation and co-occurrence, a topic brought under increasing scrutiny by the assumptions employed in metagenomic profiling tools [61]. Therefore, efficiently finding and presenting evidence for a particular trait’s presence in taxa – the essential tasks the PTBT is designed to do - serve important roles in microbial ecology, and our work provides an initial demonstration of how trait organization schemes may be designed and could inform evidence-based exploration of complex trait distributions across phylogenetically diverse and poorly characterized microbial communities.

## Materials and Methods

### Compilation of genomic data

62,291 representative genomes from GTDB release 207 (https://data.gtdb.ecogenomic.org/releases/release207/207.0/) were downloaded for annotation with six traits, as described below. We restricted our analyses only to those genomes from phyla with 100 or more representatives. To ensure that maximum estimated growth rate could be included for all genomes, only those containing at least one ribosomal protein were included in the analyses. The remaining 50,745 genomes were used for subsequent analyses. Analyses and figures were generated using Python v3.10.10. Details on the 31 phyla represented and the number of genomes per phylum are provided in Supplemental Figure 1.

### GC content and genome size

GC content and genome size were drawn directly from GTDB genome statistics, as calculated by CheckM [62].

### Oxygen tolerance

A list of NCBI taxon IDs in BacDive (accessed 6/9/2022) labeled as either ‘aerobe’ or ‘anaerobe’ were used to construct an oxygen tolerance dataset. Seven original BacDive oxygen tolerance labels from 6,629 genomes were grouped into two categories using the following scheme: anaerobe (22.2%), obligate anaerobe (1.5%), microaerophile (10.6%), facultative aerobe (0.8%), aerotolerant (0.08%), and microaerotolerant (0.03%) were assigned to ‘anaerobe’. Aerobe (46.1%), obligate aerobe (2.2%), and facultative anaerobe (8.15%) were assigned to ‘aerobe’. Each NCBI Taxon ID was matched to a representative GenBank assembly and corresponding GTDB representative genome. If no GenBank assembly was labeled as representative, we selected the highest-quality genome based on GTDB’s contamination and completeness estimates and used its GTDB species representative. When GTDB species clusters contained multiple oxygen annotations, the most common label for each cluster was paired to the respective genome. Statistics of the assembled dataset are available in the code resources (see Data Availability). 6,629 genomes with labels were identified, 662 used in final testing and 5,964 in cross-validation procedures.

A random forest model implemented in sklearn v1.0.2 was used to predict oxygen tolerance. Protein families from Pfam present in each genome were used as predictors with oxygen tolerance categories (from BacDive, Reimer et al., 2019) used as the target variable. The predicted coding sequences from representative genomes were annotated using hmmscan (HMMER v.3.3.2) and PFam release 35.0 (pfam.xfam.org) on open reading frames (ORFs) identified with Prodigal v2.6.3 [63], filtering to hits with bitscores better less than the respective Pfam entry’s “trusted cutoff.” After all matching Pfam families per gene were determined, a presence/absence table was constructed with all 62,291 genomes in release 207 (later filtered to 50,745, as above) and 17,422 (out of 19,632) observed Pfam entries. A bootstrapped logistic regression with an L1 optimizer implemented in sklearn v1.0.2 [64] was used with 100 replicates and a lambda of 1 to reduce the number of genes in the input data. The fraction of bootstraps with non-zero regression coefficients for each gene was included in later hyperparameter tuning for random forests.

We withheld 10% of data for model testing (herafter “testing data”) and retained 90% (5,964) of the 6,629 genomes in the training set for hyperparameter tuning using nested cross-validation, with 10-fold cross-validation for models and 5-fold cross-validation for combinations of parameters. We tuned the following parameters: 1) number of trees, 2) maximum tree depth, 3) minimum samples per leaf, 4) minimum samples per split, 5) bootstrapping, 6) number of features used to train each tree, and 7) number of bootstrapped logistic L1 regressions with a nonzero coefficient. Maximum training accuracy was achieved using Pfam entries with non-zero coefficients in 50-70% of randomized logistic regressions and more than 75 trees, but otherwise models were not sensitive to parameter choice. The final model used the following parameters to reduce risk of overfitting: Pfam entries present in 70% of bootstrapped logistic regressions, n_estimators = 3000, max_features = sqrt, max_depth = 13, min_samples_split=2, min_samples_leaf = 1, bootstrap = False. The random forest model achieved 92% cross-validation training accuracy and 91.4% test accuracy with modest differences in performance for aerobes and anaerobes (Supplementary Figure 4), likely due to unbalanced training data and better predictive power for aerobe-associated enzymes [28]. The model successfully recapitulated the original BacDive labels in the classification probability space (Supplementary Figure 5) and used enzymes associated with oxygen-dependent and independent metabolisms (Supplementary Figure 6).

### Inferring chlorophototrophy

Due to difficulties distinguishing rhodopsin from bacteriorhodopsin, we have focused on chlorophototrophic organisms in this study. To identify proteins and pathways which indicate phototrophy, we identified the following chlorophototrophy-related GO terms: GO:0015995, GO:1902326, GO:0036068, GO:00333005, GO:0030494, GO:0010380, GO:1902325, GO:0036067, GO:0015979, GO:0019684, GO:0019685, GO:0010109, GO:1905157, GO:0009521, GO:1905156, GO:0034357. These GO terms were used to identify 48 Pfam families of photosynthesis-related proteins, listed in Supplementary Table 3. The GenBank assembly accessions for known chlorophototrophs from Thiel et al. (2018) were used to identify which phototrophy-related Pfam families were common among phototrophic taxa. Among 698 GTDB representative genomes identified as phototrophs, 24 had none of the Pfam families. The Pfams for Photosynthetic reaction center protein (PF00124), magnesium-protoporphyrin IX methyltansferase (PF07109), and proto-chlorophyllide reductase (PF08369) were the most prevalent, indicating the bacteriochlorophyll biosynthetic mechanism can be used to identify putative phototrophs. 95.7% of the 698 genomes in Thiel et al. 2018 contained at least one complete (100% GapSeq completeness) biosynthetic process for chlorophyll a. Genomes were pre-screened using photosynthesis-related Pfam families: 12,906 of the 62,291 GTDB representative contained at least one. These 12,906 genomes were scanned with Gapseq v1.2 [65] for chlorophyll photosynthesis capacity. Any genome with at least one chlorophyll-biosynthesis-related process that was more than 90% complete was considered phototrophic for downstream analyses.

### Inferring carbon fixation

Carbon fixation potential was determined by analyzing genomes for genes associated with known carbon fixation pathways from previous literature and existing databases (Caspi et al., 2020; Kanehisa & Goto, 2000; Momper et al., 2017). Carbon fixation pathways such as the reductive tricarboxylic acid (rTCA) cycle can be difficult to distinguish from their oxidative variants using pathway completeness, so we first screened genomes for key enzymes using the HMMs from Asplund-Samuelsson and Hudson, 2021. After pre-screening, 10,166 genomes were analyzed with GapSeq to measure pathway completeness of the following MetaCyc pathways: CALVIN-PWY / Calvin Cycle, CODH-PWY / rAcoA homoacetogenic, PWY-7784 / rAcoA methanogenic, P23-PWY / rTCA I, PWY-5392 / rTCA II, PWY-5789 / 3HP/4HB, PWY-5743 / 3HP bicycle. Based on Asplund-Samuelson et al. 2021, the following completeness thresholds were used to identify putative autotrophs: CALVIN-PWY 90, CODH-PWY 95, PWY-7784 90, P23-PWY 90, PWY-5392 80, PWY-5789 90, PWY-5743 82. Using these cutoffs, 6,511 genomes out of 50,745 were identified with sufficient completeness of carbon fixation initially; only the 4,627 genomes containing Calvin Cycle or 3HP bi-cycle were considered autotrophic for the construction of the periodic table because other pathways were found in organisms not reported in the literature to be capable of carbon fixation.

### Maximum potential growth rate estimates

To estimate maximum predicted doubling times for each genome, ribosomal proteins were annotated and processed using gRodon v2 [41], a codon usage bias (CUB) based method for estimating microbial doubling time. Ribosomal proteins were used as the highly expressed gene set (from which gRodon estimates growth rates) for each genome. To annotate ribosomal proteins, we used BLASTP v2.5.0 [66] to align predicted ORFs against the growthpred database of microbial ribosomal proteins (Vieira-Silva & Rocha, 2010). ORFs with at least 50% coverage and an e-value of 1e-5 were labeled as ribosomal. 3,053 genomes had no hits. Of the remaining 50,771 genomes, 29,642 had fewer than 10 annotated ribosomal genes (Supplementary Figure 7) and taxa with lower growth rates were biased toward isolate (vs. metagenome-assembled genome) sources.

### Wavelet projection and construction of the periodic table

All subsequent analyses and figures were generated using Python v3.10.10 and ETE toolkit v3.1.2 (https://github.com/etetoolkit/ete). The GTDB r207 phylogeny was pruned using the ete3 “prune” function with keep_branch_lengths=True to contain quality-controlled species with estimates for all six traits (50,745 total). A Haar-like wavelet basis was computed for this tree with code from Gorman and Lladser, 2022 (https://github.com/edgor17/Sparsify_Ultrametric) and used to compute a 50,744 × 6 matrix of Harr-like wavelets coefficients per trait for each non-root interior node. The vector of Haar-like wavelet coefficients for each trait were divided by the L2 norm, yielding a normalized vector of variances explained per wavelet coefficient/ internal node. To identify nodes which would define a periodic table composed of functionally uniform groups, the normalized wavelet coefficients were sorted and cumulatively summed starting with the largest wavelet until a selected amount of variance was captured (60%) for each trait. Subtrees that did not contain any of the 72 total nodes identified by this procedure were collapsed using a custom tree-pruning algorithm.

Algorithm to construct a collapsed tree *T*_*W*_:

1. Define a set of nodes *W* to avoid collapsing
2. Begin with the original phylogeny *T*
3. Traverse *T* Starting with the root of *T*;

For node *N* in *T:*

a. If *N* is a leaf of *T*:
  i. mark it as a leaf of *T*_*W*_.
b. Else if all descendants of *N* are not in *W*:
  i. mark *N* as a leaf of *T*_*W*_.
c. Else:
  i. Continue
d. Build *T*_*W*_using only marked nodes as leaves; collapse all unmarked subtrees.

This algorithm retains nodes only if they belonged to a wavelet clade or were necessary to maintain the tree structure connecting retained nodes. This algorithm pruned the original 50,745 tips to 272 tips (clades from the original phylogeny). Re-computing the trait data from the wavelet coefficients in this pruned tree would create a trait distribution that captured 60% or more of the variation from the original data. These 272 clades were used to draw the periodic table. A 272 × 272 patristic distance matrix of these clades and a 272 × 6 matrix of the scaled median trait values for the leaves associated with each clade was used to compute a three-dimensional ENS-t-SNE embedding with 4000 iterations and a perplexity of 50 using mview (https://github.com/enggiqbal/MPSE-TSNE). The images from the resulting embedding were combined and points were snapped to a grid using the Hungarian algorithm implemented in sklearn. The grid layout, median trait values for each clade, and phylogenetic tree were used to generate the periodic table diagram and tree diagrams in Figures 4 and 5 using custom Python scripts and the seaborn [67], numpy [68] and matplotlib [69] libraries.

## Acknowledgements

We want to thank JL Weissman and members of the Fierer Lab for their insights and willingness to engage in productive discussions about this work. We also want to thank the Information Technology team at CIRES for computational support. We are deeply indebted to the team responsible for the Genome Taxonomy Database (GTDB) which made this work possible.

## Author contributions

M.H. and N.F. conceived this study with all analyses conducted by M.H. with E.G. and M.L. helping with the Haar-like wavelet analyses. Writing was led by M.H. and N.F. with reviewing and editing of the manuscript by E.G. and M.L. All authors read and approved the final manuscript.

## Conflicts of interest

The authors declare no conflicts of interest.

## Funding

This work was supported by grants from the US National Science Foundation (DEB 2126106, AW5809-826664, OPP 2133684) and funding provided to the Center for Microbial Exploration by the University of Colorado Boulder.

## Data Availability

Intermediate data files and code used to generate figures and analyses are available on GitHub: https://github.com/realmichaelhoffert/bacterial_periodic_table/tree/main

## Notes

### Competing Interest Statement

The authors have declared no competing interest.

