## Supplementary Information and Figures for "A periodic table of bacteria?: Mapping bacterial diversity in trait space"

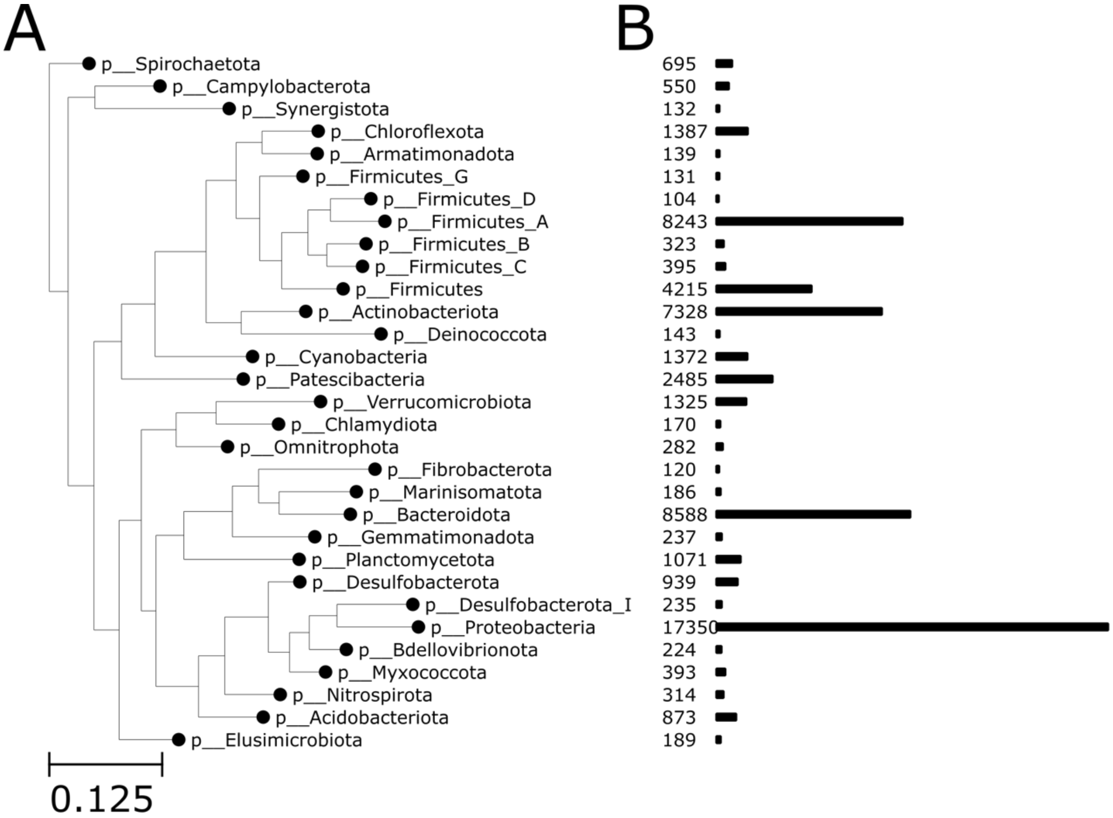


**SF1** – (A) Reference GTDB phylogeny of 31 bacterial phyla with at least 100 genomes used in this study. (B) Bars and values showing the number of representative GTDB genomes per phylum.

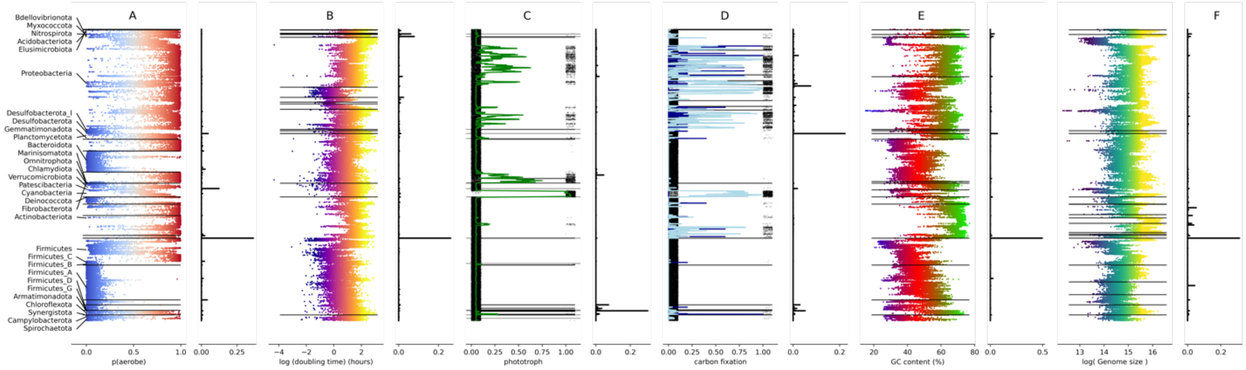


**SF2** – Panels A-F show distributions of L2-normalized Haar-like wavelet coefficients for each trait: (A) oxygen tolerance (B) maximum potential growth rate (C) carbon fixation capacity (D) phototrophy (E) GC content and (F) genome size. Panels A-F contain trait values for each genome in postorder in the left plot, and normalized Haar-like wavelet coefficients placed at the y-axis location of the internal node they correspond to in the right plot. Each value in the vector of wavelet coefficients for a trait corresponds to an internal node of the phylogeny; it

describes the amount of variance in a trait uniquely attributed to the differences between the left subtree and the right subtree of the node. Each coefficient is normalized to represent the fraction of the trait's total variance it describes; in a tree with exactly 50% of the species with a trait, and 50% without it, a single coefficient with a magnitude of 1 would be found at the common ancestor of the positive and negative groups. Nodes with larger coefficients have left and right sub-trees that are more different with respect to each trait than nodes with smaller coefficients. Additional lines are drawn over the trait data to mark the edges of the left and right subtrees for the clades with the ten largest wavelet coefficients, illustrating how the wavelet coefficients reflect trait shifts in the data.

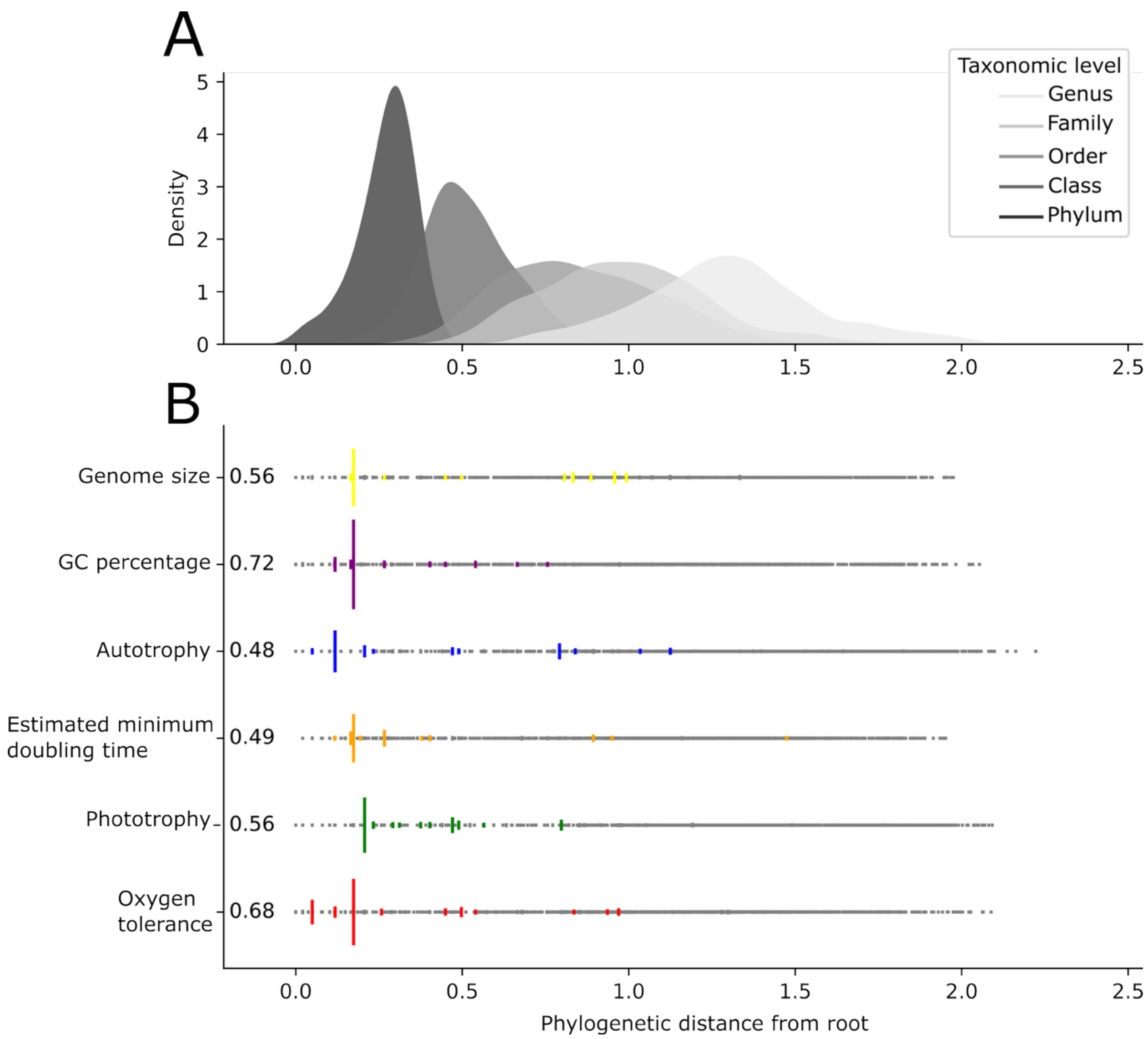


**SF3 –** The average phylogenetic depth of ancestral nodes (A) and nodes with significant trait variation (B) for each of the six traits, based on the Haar-like wavelet analyses (see Methods). Panel A shows the distribution of root-node distances for the ancestral nodes of each taxonomic level, with shorter distances from the root corresponding generally to more ancient phylogenetic splits and higher taxonomic levels. (B) The phylogenetic distance from the root versus magnitude of Haar-like wavelet coefficients, a measurement of the amount of trait variance associated with a particular node. The ten largest wavelets are colored for each trait: the total variance captured by the ten largest coefficients is labeled along the y-axis for each trait.

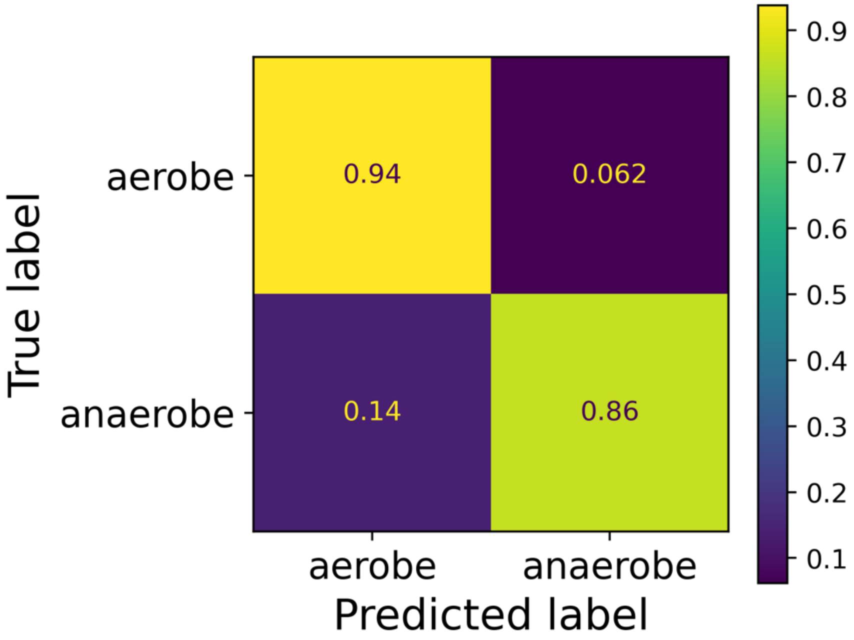


**SF4** – Confusion matrix for the oxygen tolerance predicting random forest model evaluated on the withheld test dataset (n=664). The matrix shows the fraction of true and false predictions for true members of each class, indicating the total percent of correct (upper-left, bottom-right) or incorrect (bottom-left, upper-right) classifications across classes.

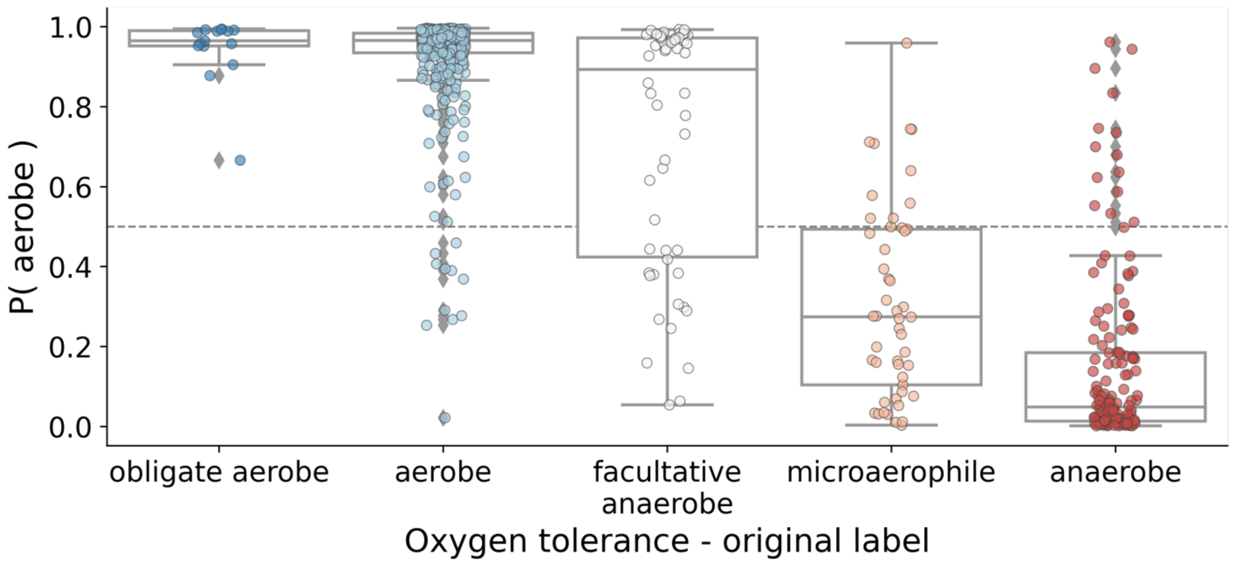


**SF5** – Comparison of the random forest's prediction probability versus the original BacDive oxygen tolerance category for 664 bacteria in the test dataset. The y-axis shows the predicted probability of being oxygen tolerant, or probability of classification as "aerobe," produced by the model. The x-axis labels each bacteria using the original BacDive categories, which were grouped into "aerobe" or "anaerobe" based on the class they were most similar to. This plot illustrates that the model can successfully distinguish intermediate degrees of oxygen tolerance.

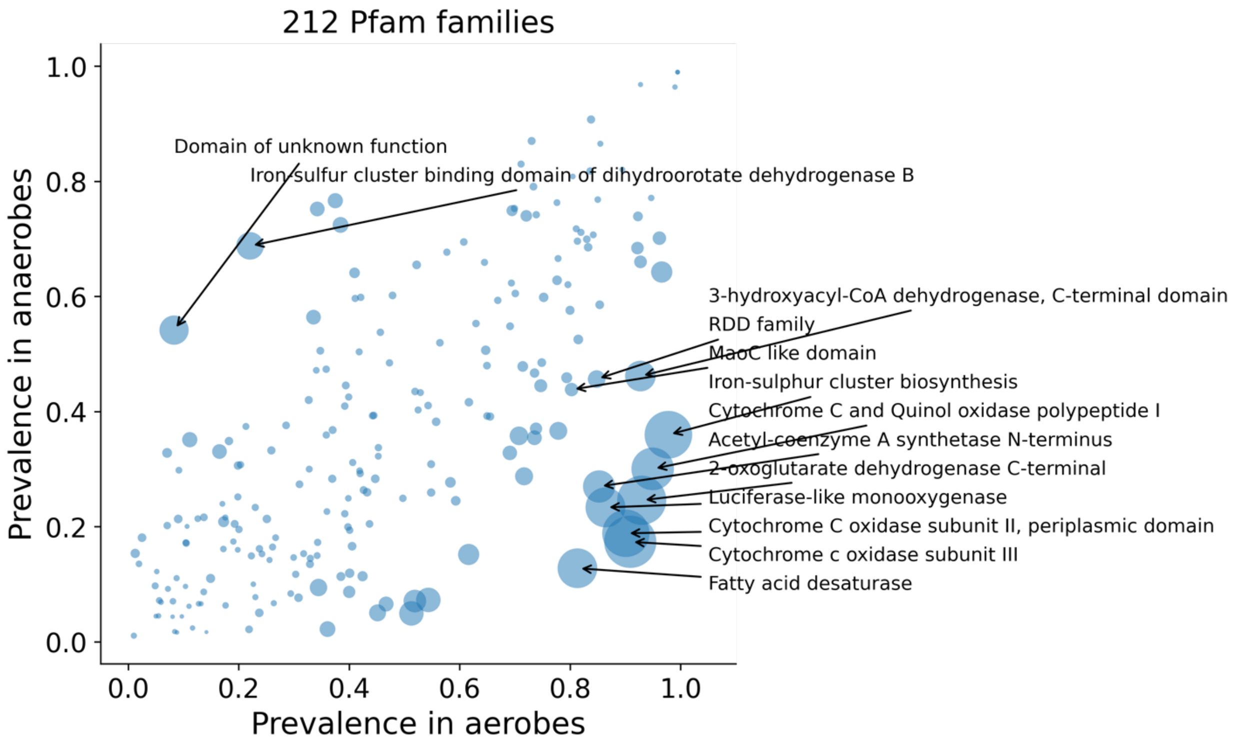

**SF6** – Difference in prevalence of 212 Pfam families used by the random forest to predict oxygen tolerance in aerobic bacteria (x-axis) and anaerobic bacteria (y-axis) from the training dataset. Points are sized by random forest feature importance to illustrate that points with larger differences in prevalence are more important to distinguishing aerobes from anaerobes. High-importance genes are annotated with descriptions from the Pfam database.

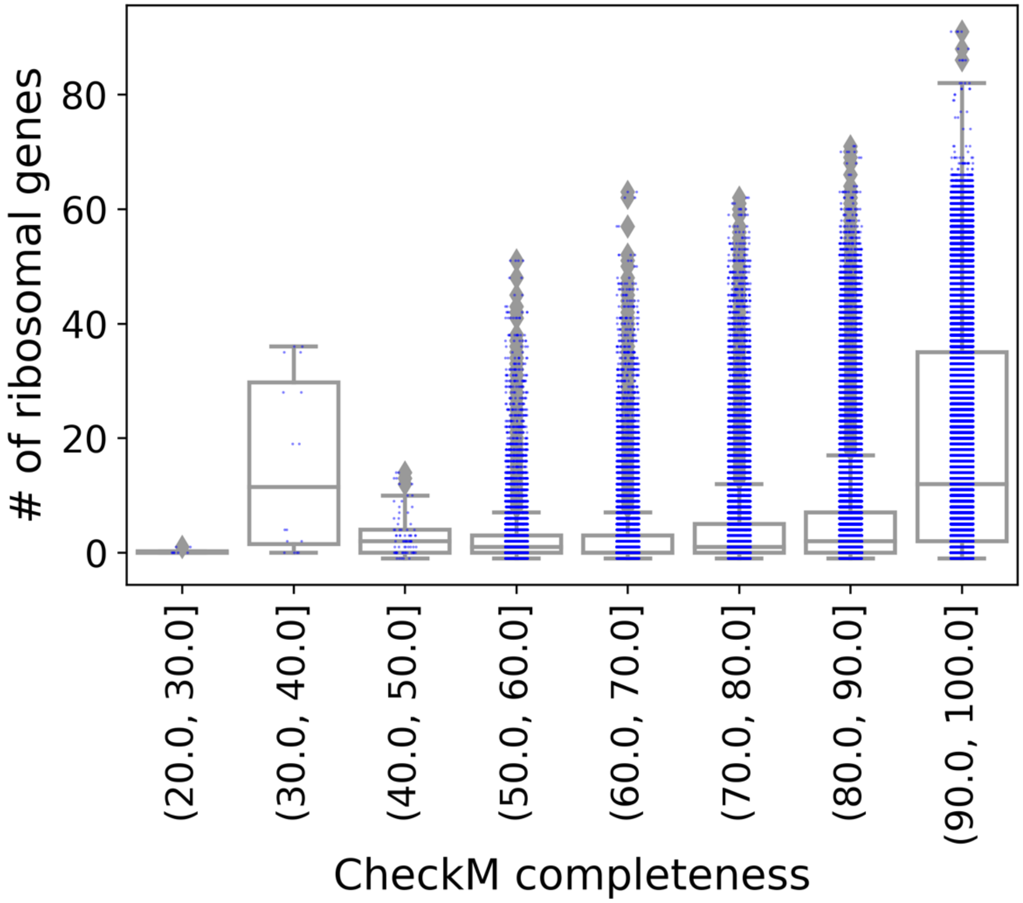


**SF7** - Number of ribosomal proteins versus genome completeness for all v207 GTDB representative genomes showing that the number of ribosomal proteins detected decreases as genome completeness is reduced. Genome completeness likely influences the accuracy of minimum doubling time estimates produced by gRodon.

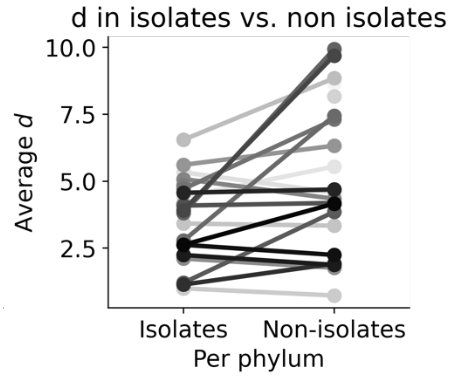


**SF8** – Average predicted minimal doubling times from gRodon in isolates and non-isolates for the 31 largest bacterial phyla shows substantial bias in growth rate for cultured species. Lines connect pairs of measurements for each phylum and are colored by the number of observations in the phylum.
